# An automated platform for structural analysis of membrane proteins through serial crystallography

**DOI:** 10.1101/2021.06.03.446146

**Authors:** Robert D. Healey, Shibom Basu, Anne-Sophie Humm, Cedric Leyrat, Xiaojing Cong, Jérome Golebiowski, Florine Dupeux, Andrea Pica, Sébastien Granier, José Antonio Márquez

## Abstract

Membrane proteins are central to many pathophysiological processes yet remain very difficult to analyze at a structural level. Moreover, high-throughput structure-based drug discovery has not yet been exploited for membrane proteins due to lack of automation. Here, we present a facile and versatile platform for *in meso* membrane protein crystallization, enabling rapid atomic structure determination at both cryogenic and room temperature and in a single support. We apply this approach to two human integral membrane proteins, which allowed us to capture different conformational states of intramembrane enzyme-product complexes and analyze the structural dynamics of the ADIPOR2 integral membrane protein. Finally, we demonstrate an automated pipeline combining high-throughput microcrystal soaking, automated laser-based harvesting and serial crystallography enabling screening of small molecule libraries with membrane protein crystals grown *in meso.* This approach brings badly needed automation for this important class of drug targets and enables high-throughput structure-based ligand discovery with membrane proteins.

**Highlights:**

- A fully automated, online workflow enables rapid determination of membrane protein structures by serial X-ray crystallography (SSX).
- High resolution room temperature and cryogenic structures of ADIPOR2 provide insights into the dynamic nature of receptor:ligand interactions.
- A web-based application allows remote user-guided experimental design and execution.
- An automated SSX-based ligand discovery pipeline for integral membrane proteins is introduced.

## Introduction

Membrane proteins are key regulators of most physiological processes and represent attractive targets for drug discovery programs (Andrews et al., 2014; Congreve et al., 2020). Their structure determination remains a powerful strategy to gain insight into their function and accelerates drug development (Andrews et al., 2014; Ishchenko et al., 2019). One of the most successful methods to obtain high-resolution structures from membrane proteins relies on *in meso* crystallization (i.e. lipidic cubic phase, LCP) and X-ray diffraction (Caffrey, 2015; Landau and Rosenbusch, 1996; Landau et al., 1997; Liu and Cherezov, 2011; Pebay-Peyroula, 1997). This strategy has been particularly successful at solving structures of the G protein-coupled receptor (GPCR) family and other integral membrane proteins (Caffrey, 2015; Rasmussen et al., 2011; Vasiliauskaité-Brooks et al., 2017, 2018). *In meso* crystallization remains a long process with challenges at every step. This may justify the under-representation of membrane protein structures in the Protein Data Bank (wwwPDB)(Berman, 2000). Structure-based ligand discovery efforts targeting membrane proteins, such as fragment screening, remain inadequate due to technical barriers in experimental set-ups.

*In meso* crystallization trials are typically performed with sandwich plates, where the sample mesophase is squeezed between two glass plates (Caffrey and Cherezov, 2009; Cherezov et al., 2004). Membrane protein crystals grown *in meso* are often of micrometer-sized dimensions and decay rapidly due to radiation damage. Thus, analysis by X-ray diffraction is usually performed through the so-called serial synchrotron crystallography (SSX) approach, applied both at ambient and cryogenic temperatures (Gati et al., 2014; Huang et al., 2015, 2016; Weierstall et al., 2014). For room temperature experiments, delivery of LCP membrane protein crystals to an X-ray beam for SSX can be achieved using sample jets (Botha et al., 2015; Nogly et al., 2015, 2018; Weierstall et al., 2014; Weinert et al., 2017). The LCP injection method has many benefits for time-resolved experiments. However, it requires large quantities of protein, extensive optimization of sample preparation, manual delivery protocols, and long times for data collection at the beamline. For cryogenic SSX analysis, crystals grown in sandwich plates are manually harvested – for which one of the glass plates must be removed, or broken, and the crystals manually recovered and flash-frozen in a matter of seconds before the crystals are damaged (Caffrey and Cherezov, 2009; Li et al., 2012). Even in experienced laboratories, manual LCP crystal harvesting is a difficult, time-consuming and inefficient process. Post-crystallization treatments such as ligand or fragment soaking in an automated and straightforward manner are not possible. A promising approach to allow soaking of *in meso* crystals has been suggested in which crystallization occurs between two plastic films (Huang et al., 2015, 2016). However, this method poses technical challenges, entailing manual handling and low throughput (Huang et al., 2018). Currently, the field lacks automation that could support higher-throughput approaches (Rucktooa et al., 2018), a necessity to expedite structure-based ligand discovery for membrane proteins.

The CrystalDirect method was initially developed to enable automated recovery of crystals from soluble proteins grown through vapor diffusion (Cipriani et al., 2012; Márquez and Cipriani, 2014; Zander et al., 2016). In this approach, crystals are grown on thin films in a commercially available 96-well crystallization plate (a CrystalDirect plate, **Figure 1**) and harvested directly on X-ray data collection pins by a robotic laser-based excision system (Zander et al., 2016). This approach has thus far been applied to processing crystals from soluble proteins (Aragón et al., 2019; Bezerra et al., 2017; Martin-Malpartida et al., 2017). However, its application to membrane proteins was limited by incompatibility of standard LCP sandwich plates with the CrystalDirect laser-excision technology. To the best of our knowledge, automated harvesting of crystals grown *in meso* has not been described to date and manually harvesting crystals remains a major technical challenge, limiting high-throughput applications.

**Figure 1.**
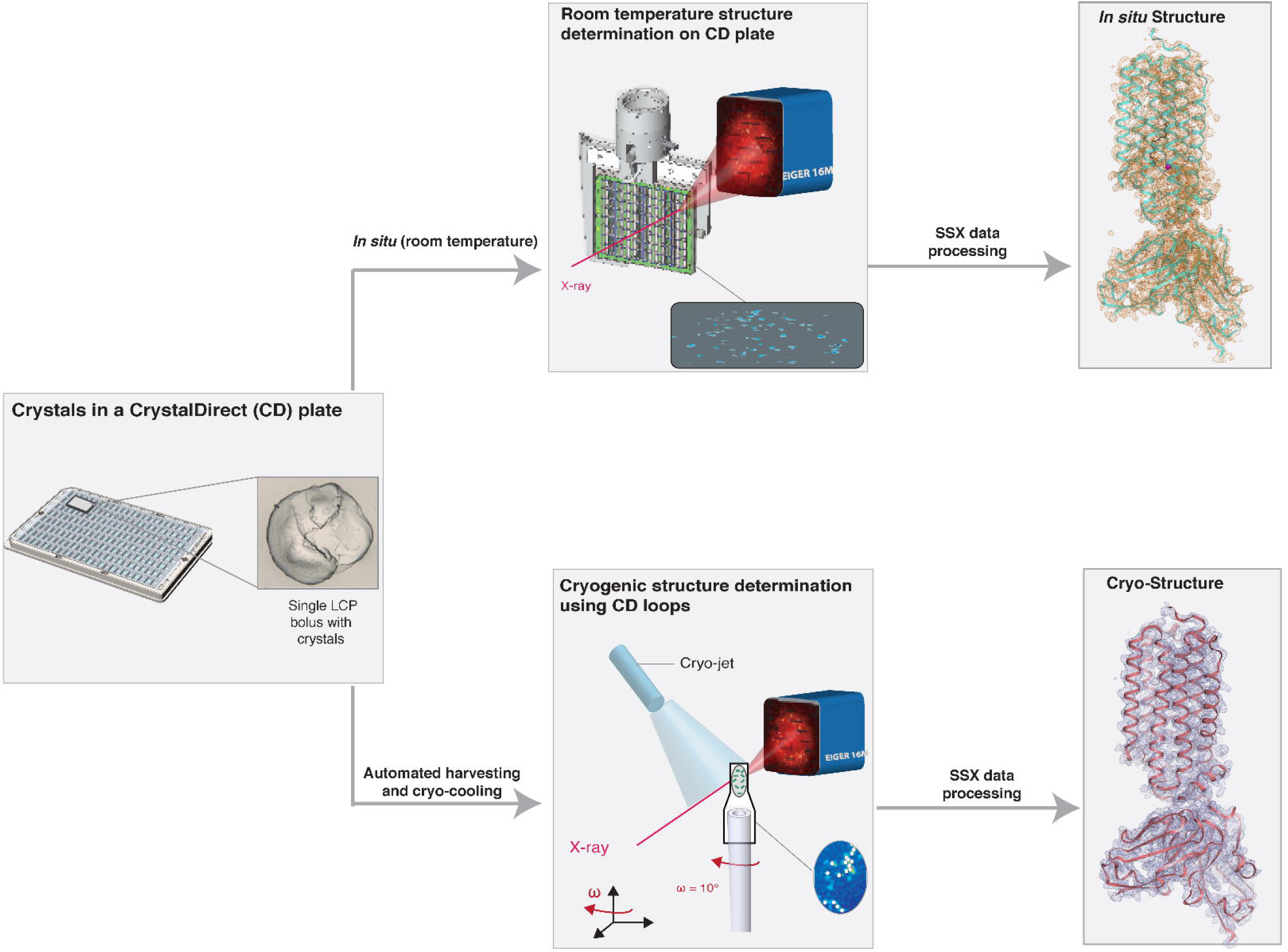
*In meso* crystallization and SSX analysis of membrane proteins in CrystalDirect plates. The **left-panel** shows a CrystalDirect (CD) plate with LCP crystallization experiments (**inset**). Once LCP crystals have grown, two different workflows for structural analysis can be applied under the same crystallization conditions. Room temperature *in situ* X-ray diffraction data can be collected directly from the CrystalDirect plate (**Middle-panel top**). For cryogenic structures (**Middle-panel bottom**) a workflow including automated laser-based harvesting and cryo-cooling can be utilized. Room temperature and cryogenic structures of ADIPOR2 (left, top and bottom respectively) obtained with this approach are shown.

Here, we introduce a platform to grow and monitor membrane protein crystals in CrystalDirect plates using two pharmacologically important integral membrane protein targets – human alkaline ceramidase 3 (ACER3) and adiponectin receptor 2 (ADIPOR2). Using CrystalDirect as a support for LCP crystallization allows SSX diffraction to be monitored *in situ* at ambient and cryogenic temperatures, enabling the direct comparison of ADIPOR2 structures solved at different temperature and revealing insights into ADIPOR2-lipid interactions and conformational dynamics. Moreover, we demonstrate how this technology can be applied to high throughput ligand screening with membrane proteins. To facilitate automated experimental workflows, we have developed the web-based Crystallographic Information Management System (CRIMS). This information system allows automated tracking of crystallographic experiments and web-based access to experimental design and evaluation tools encompassing protein sample registration, crystallization set-up and monitoring, crystal harvesting as well as shipment to and recovery of processed data from synchrotrons. Through the combination of the CystalDirect technology and CRIMS software the HTX facility in Grenoble provides access to a resource for the rapid structural analysis of membrane proteins *in meso,* removing major hurdles and allowing rapid structural analysis of membrane proteins. This resource is open to scientists worldwide through different access programs and could be replicated elsewhere.

## Results and Discussion

### Automated harvesting of membrane protein crystals

We have developed a new method for LCP crystallization compatible with the CrystalDirect technology, thereby enabling automated harvesting, soaking and cryo-cooling of crystals grown in meso (**STAR Methods**). In this method, LCP crystallization experiments are set inside the vapour diffusion chambers of a CrystalDirect plate, instead of on a sandwich plate. In order to avoid evaporation, the reservoir side of the vapour diffusion well is filled with an excess volume of crystallization solution and the plate sealed from the top with a standard film (**Figure 1, Figure 3**). Under these conditions, experiments are stable for several weeks. Once crystals develop, the plates are transferred to the CrystalDirect harvester robot, where film pieces containing the crystal-laden LCP bolus or parts of it are automatically excised, mounted onto a standard data collection pin and transferred to a cryogenic sample storage device (**Figure S1**).

The overall management of crystallization experiments presented here is made possible through the Crystallographic Information Management System (CRIMS), a web-application for the design, evaluation and analysis of crystallographic experiments (https://htxlab.embl.fr/). Using CRIMS, scientists can register protein samples, design crystallization screens, monitor crystal growth and define automated harvesting (**Figure S2**). Once crystals have been identified, harvesting is performed by the CrystalDirect harvester robot (**Figure S2**). The CrystalDirect robot then transfers the sample pin to a standard sample holder in a cryogenic storage device, while CRIMS records sample locations. The entire process is outlined in **Supplemental Video 1**. A single harvesting operation takes approximately one minute and up to 312 samples (pins) can be prepared in a single session.

For this study we chose the seven-transmembrane protein, adiponectin receptor 2 (ADIPOR2), a receptor activated by protein hormone adiponectin. Adiponectin exerts a pleiotropic action in mammals and is implicated in metabolic and cardiovascular related disorders (Straub and Scherer, 2019), small molecules replicating adiponectin effects have long been sought after for clinical applications in the treatment of metabolic and cardiovascular related diseases (Okada-Iwabu et al., 2013). The CrytalDirect LCP method produced crystals under the same precipitant conditions and with similar size and morphology of those previously reported using the classic sandwich glass plate approach (Vasiliauskaité-Brooks et al., 2017). We collected cryogenic SSX diffraction data on ADIPOR2 crystals harvested from single *in meso* boluses. Loops containing multiple ADIPOR2 microcrystals were defined in CRIMS and harvested by the CrystalDirect harvester such that a single loop contained 10-100 crystals (**Figure S3**). Diffraction data was collected at the Swiss Light Source (SLS) on PXI (X06SA) from each microcrystal as 10° wedges of minisets as previously described (Basu et al., 2019; Wojdyla et al., 2016). A complete ADIPOR2 dataset was obtained from a single harvested loop, requiring 300 ng of protein. The structure of ADIPOR2 harvested automatically from a CrystalDirect plate was resolved at 2.4 Å (**Figure 1**); this resolution is equivalent to that previously reported by manual harvesting from glass plates (Vasiliauskaité-Brooks et al., 2017) indicating our automated approach does not lead to any reduction in data quality.

We also crystallized an intramembrane sphingolipid hydrolase, alkaline ceramidase 3 (ACER3), using this method. In glass sandwich plates, ACER3 crystals tend to grow in the sponge phase and are particularly challenging to harvest manually. Using the CrystalDirect harvester, all attempted ACER3 crystal harvests were successful. Nine harvested loops were required to collect a complete dataset for ACER3 employing the same SSX methodology as for ADIPOR2. The data collection required 36 minutes of synchrotron beam time at the Swiss Light Source. A high-quality structure of ACER3 at 2.6 Å resolution was resolved, requiring 1.6 μg of protein (**Figure S4**). This resolution was equivalent to that previously reported using classic sandwich glass plates (Vasiliauskaité-Brooks et al., 2018), demonstrating that *in meso* crystallization in CrystalDirect plates does not compromise data quality.

### *In situ* membrane protein serial crystallography

The CrystalDirect plates can also be directly used for in situ X-ray experiments (**Figure 1**). We used this approach for room temperature analysis of ADIPOR2 through in situ diffraction (**STAR Methods**). We collected *in situ* datasets from ADIPOR2 crystals by mounting the plate directly on an SBS-plate goniometer at the P14 beam line of the Petra III synchrotron (Hamburg, Germany). The crystals were exposed to the X-ray beam and diffraction recorded at room temperature with a continuously oscillating raster-scan across each LCP bolus (Gati et al., 2014) (**Supplemental Video 2**). A collection strategy that optimised beam transmission and exposure time whilst minimising radiation damage was applied (**STAR Methods**). A complete dataset was acquired from five boluses in different wells in approximately 20 minutes of beamtime representing a total of 1.5 μg of protein. At room temperature, ADIPOR2 crystals diffracted to 2.9 Å resolution (**Figure 1**). An added benefit of using the CrystalDirect plate for *in meso* crystallization is that structural analysis can be carried out at cryogenic and room temperature in the same support and under the exact same conditions in a straightforward manner. Unlike existing sample-to-beam delivery methods like LCP-jets or microchips (Botha et al., 2015; Nogly et al., 2015; Weierstall et al., 2014) our approach requires essentially no optimization. In CrystalDirect plates, crystal growth conditions are easily translated from glass sandwich plates. CrystalDirect plates can also be used for primary LCP crystallization screening.

### Structural analysis at different temperatures reveals dynamics of ADIPOR2 interaction with endogenous lipid

Cooling protein crystals to cryogenic temperatures allows for a reduction in radiation sensitivity, but can also decrease the heterogeneity of protein conformations (Russi et al., 2017). By collecting diffraction data at ambient and cryogenic temperatures alternative conformations relevant to protein function can be revealed (Fraser et al., 2009, 2011).

Adiponectin receptors belong to a larger family of transmembrane receptor putatively assigned with hydrolase activity due to a conserved active motif of three histidines coordinating a Zinc ion (Pei et al., 2011). The adiponectin receptors 1 and 2 (ADIPOR1/2) function as ceramide hydrolases, utilising the bioactive lipid ceramide as a substrate and producing sphingosine and free fatty acids, such as oleic acid (Vasiliauskaité-Brooks et al., 2017). The structures of ADIPOR1 and ADIPOR2 reveal an internal cavity containing the active site, enclosed by seven transmembrane helices (7TMs). The mechanism by which substrate enters the cavity to be processed into products is not yet characterized. Recent structural data has revealed the existence of distinct conformations of transmembrane helix 5 (TM5) for ADIPOR1 and ADIPOR2, so called open and closed, respectively (Tanabe et al., 2020; Vasiliauskaité-Brooks et al., 2017, 2019).

In the ADIPOR2 structures determined here, TM5 is in a closed conformation and a free fatty acid product, oleic acid (OLA), is inside the catalytic cavity (**Figure 2A** and **Figure 4G, H, K**). Overall, the room temperature and cryogenic structures showed similar backbone conformations (RMSD = 0.3 Å). However, we did observe a conformational change in the bound oleic acid (OLA) molecule (**Figure 2** and **Figure 4**). To explore the relevance of this enzymatic product conformational difference, we performed replica-exchange molecular dynamics simulations of ADIPOR2 at room temperature, using OLA-depleted and OLA-bound cryogenic structures as starting models. During the OLA-bound simulation, OLA rapidly adopted a conformation corresponding to that of our room temperature structure, and initially continued to oscillate around this position (**Figure 2C**). As the OLA-bound simulation proceeded, the intracellular terminal of ADIPOR2 TM5 suddenly transitioned into a more open position, coinciding with a displacement of OLA away from the active site zinc and towards the lipid bilayer (**Figure 2B** and **Figure 2C**). This simulation captures the dynamics required for ADIPOR2 to release its enzymatic product, OLA. The open TM5 conformation observed in the MD simulations resembles that of the open structure reported in the ADIPOR1 crystal structures (Tanabe et al., 2020; Vasiliauskaité-Brooks et al., 2017), suggesting conserved conformational transition in the ADIPOR family associated to product release and necessary for enzymatic activity. In simulations performed in the absence of OLA, TM5 was observed to gradually close (**Figure 2B**). The versatility of CrystalDirect plates as *in meso* crystallization supports allowed us to capture different conformational states of intramembrane enzyme-lipid complexes. To maximize comparability, we were able to use identical crystallization conditions for both cryogenic and *in situ* experiments with protein prepared from a single purification batch, enabling a true comparison between the resulting protein structures. Our results highlight the potential of this technique, when combined with computational approaches, to analyze the structural dynamics of membrane proteins. This information is key to provide mechanistic understanding of protein function and can be exploited for drug design.

**Figure 2.**
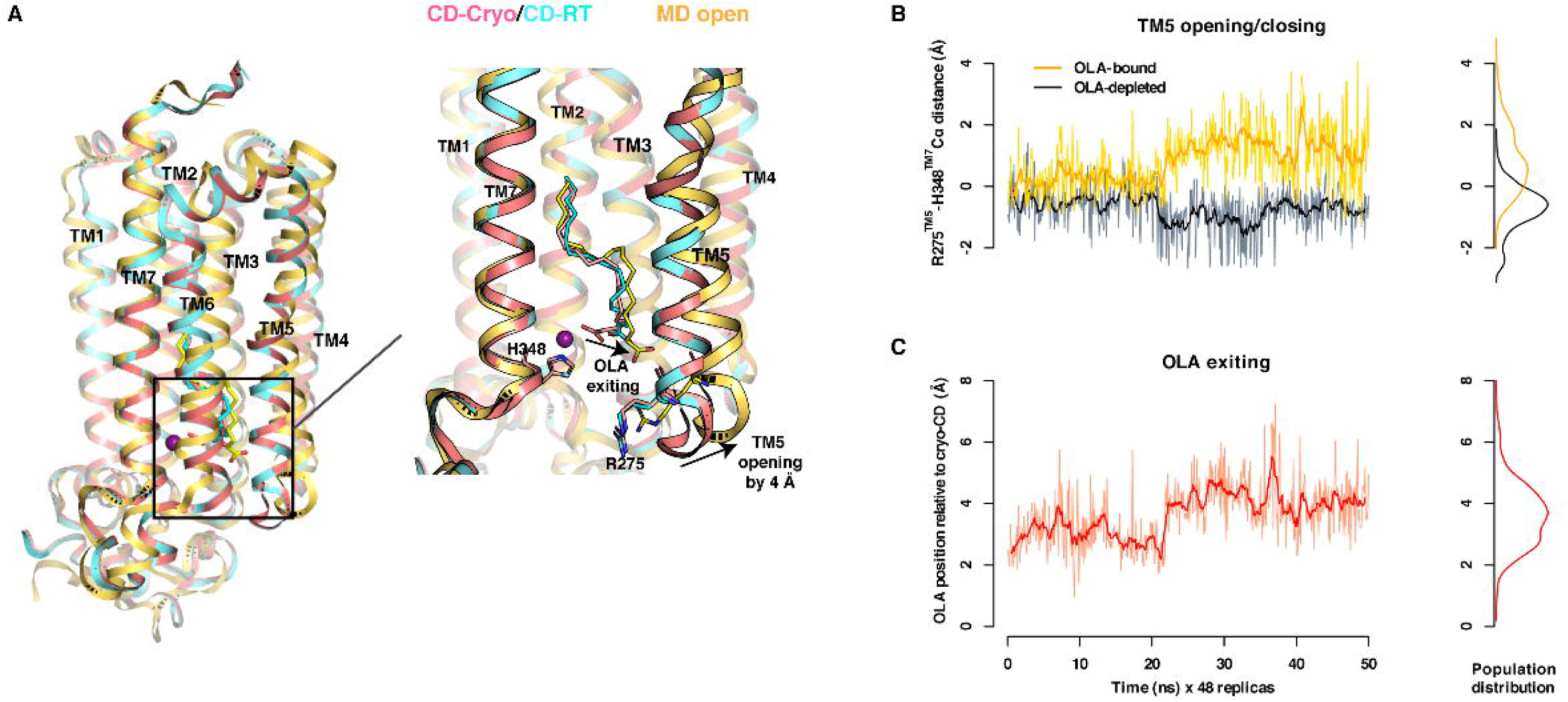
ADIPOR2-lipidic product dynamics. (**A**) Overlay of CrystalDirect structures: cryogenic (magenta) and room temperature (cyan) with MD simulation (yellow). (**B**) MD trajectory plots overlaying OLA-bound (yellow) and OLA-depleted (black) simulations, measuring ADIPOR2 transmembrane helix 5 (TM5) opening of 4 Å (distance: R275^TM5^-H348^TM7^) with density plot of the trajectory shown on the right. (**C**) MD trajectory plot of OLA-bound simulation measuring displacement of OLA from the active site by 3.9 Å (distance: catalytic Zinc atom – carboxylic head of OLA). The Spearman’s correlation coefficient between TM5 opening and OLA movement is 0.74.

### A pipeline for automated ligand soaking *in meso*

To expand the possibilities of high-throughput structure-based ligand discovery on membrane protein targets, we have developed an automated pipeline. This pipeline incorporates LCP crystal production, ligand soaking, crystal harvesting, cryogenic SSX data collection and structure determination of microcrystals grown *in meso* (**Figure 3**). Automation of crystal manipulation is a key consideration for any type of screening, as it enables high throughput and reduces the possibility of human error. Soaking of LCP crystals in traditional plates requires manual separation of one of the two sandwich layers holding the LCP bolus, which is a delicate operation and can result in sample damage. This has limited the application of high throughput ligand screening approaches for membrane proteins. The resealable CrystalDirect plate is an optimal support for *in meso* experiments, allowing direct access to experiments for post-crystallization manipulation without damaging the bolus or the crystals. To soak ADIPOR2 crystals, the top sealing film of the plate was removed, ligand solutions added with a mosquito robot and the plate was resealed for the desired soaking time (**Figure 3; top-row** and **STAR Methods**). Addition of 96 soaking solutions to an entire microplate takes < 2 min in total, indicating rapidity and high-throughput of our automated soaking pipeline. After addition of the ligands, experiments were incubated inside a crystal imaging robot and images of soaked crystals were used to define harvesting plans. Following soaking, crystals were harvested and prepared for synchrotron shipment as outlined for ADIPOR2 and ACER3 (**Figure 3; mid-row**).

**Figure 3.**
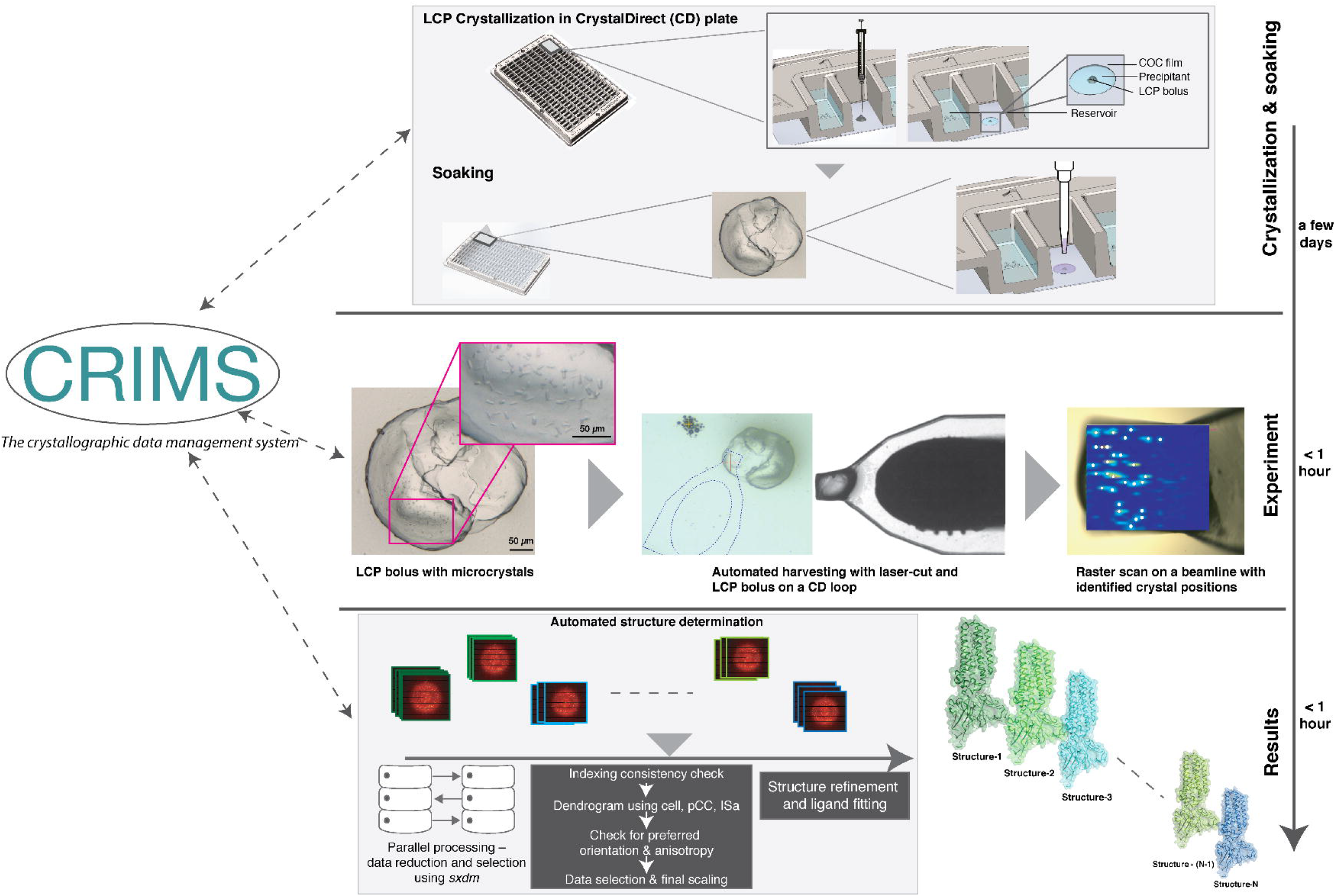
A pipeline for high throughput ligand discovery with membrane protein crystals. Following LCP crystallization in CrystalDirect plates, crystals can be directly accessed by removal of the top plastic seal on the plate (**top row**). Soaking solutions are introduced to the experiment by a pipetting robot after which the plate is re-sealed and incubated for the necessary time **(top row)**. Following soaking, LCP crystals are *auto*-harvested with the CrystalDirect robot into pins and stored in UniPucks for shipment to synchrotron **(middle-row; Experiment)**. Serial diffraction data is collected for each uniquely soaked bolus and merged following established protocols before structure refinement and ligand fitting **(Bottom-row; Results)**. The flow of data and automation of our pipeline is maintained through the CRIMS-information management system through which users can access, monitor, and analyze data via web-interfaces.

Processing of SSX diffraction data for a large number of samples remains a challenging computational task. Efficient detection of well-diffracting microcrystals via raster scanning, and automatic triggering of multiple dataset collection with small-wedges are imperative for high-throughput SSX data collection (Basu et al., 2019). On-the-fly data processing and feedback together with efficient selection of useful datasets is also necessary to enable high-throughput screening of samples. In this work, cryogenic SSX measurements were carried out at the Swiss Light Source, where data collection and real-time data processing, selection, and merging are available through the *sxdm* package (Basu et al., 2019).

To confirm that *in meso* crystals grown in CrystalDirect plates could be soaked with small molecules, we soaked ADIPOR2 crystals with heavy metal containing organic caged molecules, typically used as phasing agents (Engilberge et al., 2017; Girard et al., 2003). Following incubation with ligand, automated harvesting was performed on regions of interest containing dozens of microcrystals. For initial soaking experiments, we soaked several *in meso* boluses for each ligand, then collected cryogenic SSX diffraction data. The electron density maps show two distinct binding sites for Tb-XO4, in clefts on the surface of the antibody fragment portion of the ADIPOR2:scFv complex (**Figure 4D, I**). The structure of ADIPOR2 soaked with Gd-DO3 revealed binding of the compound to transmembrane helix 3 (**Figure 4E, J**). Both ADIPOR2 metal complexes were solved at 3 Å resolution (**Table 1**).

**Figure 4.**
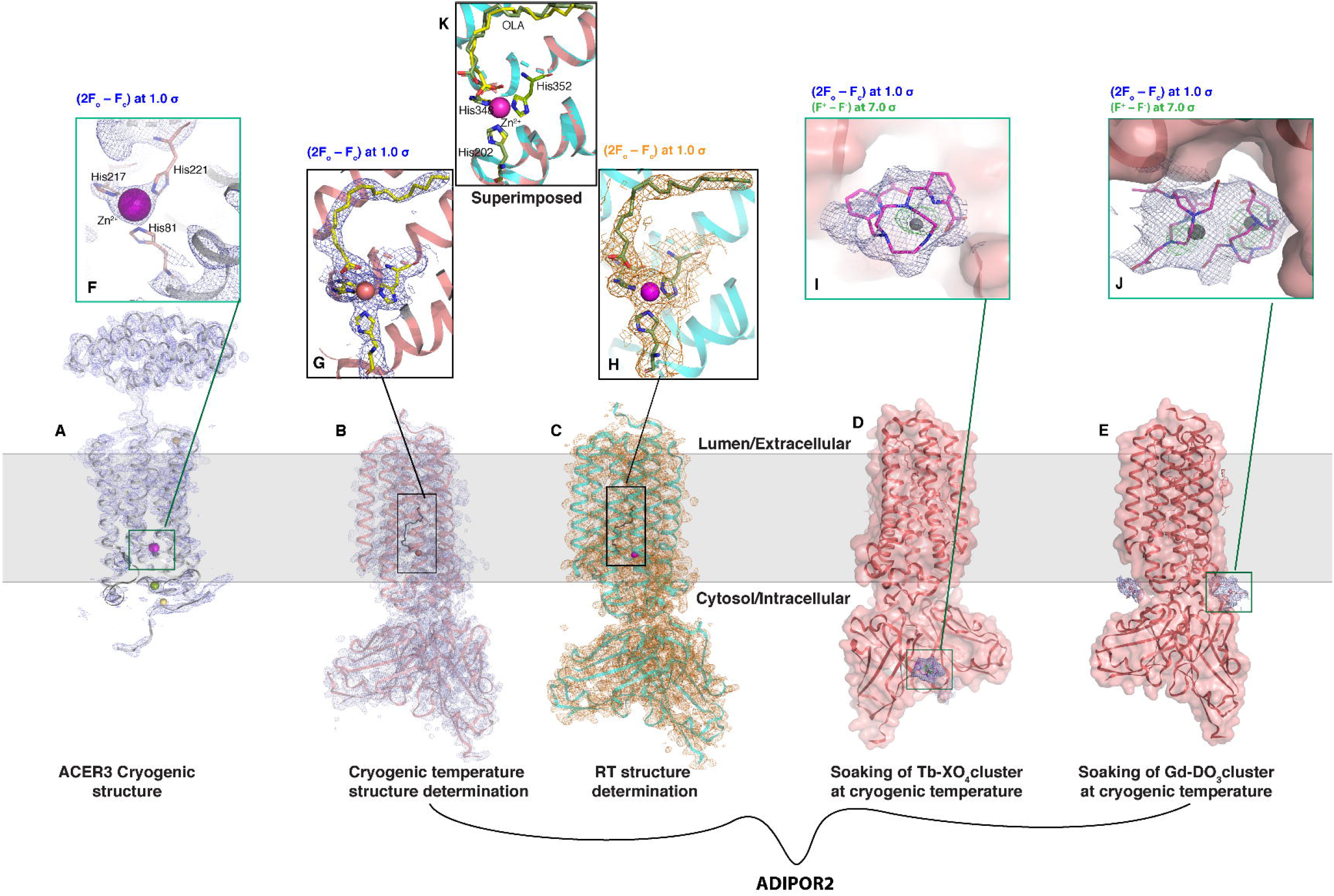
High throughput structural analysis of integral membrane proteins ADIPOR2 and ACER3. Structures of the ACER3 and ADIPOR2 seven-transmembrane enzyme receptors were obtained with the workflows presented in **Figure 1**. Protein crystal structures are shown with the membrane plane indicated as a grey zone. (**A**) ACER3 cryogenic structure, shown as cartoon in grey colour and (2F_o_ – F_c_) map in blue color, (**B**) ADIPOR2-scFv cryogenic structure, shown as cartoon in salmon color with (2F_o_ – F_c_) map (**C**) ADIPOR2-scFv room temperature structure, shown as cyan cartoon with orange (2F_o_ – F_c_) map. (**D, E**) ADIPOR2-scFv structures shown as surface structures, highlighting Tb-complex and Gd-complex as soaked clusters with (2F_o_ – F_c_) map in blue. All (2F_o_ – F_c_) maps are contoured at 1.0 sigma. (**F–H**) Inset views of active sites, showing Zn^2+^ atoms as magenta-colored spheres, coordinated by His residues shown as sticks. The respective (2F_o_ – F_c_) maps, matching the color. (**K**) superposition showing distinct conformations of the oleic acid molecule at the catalytic site of ADIPOR2 in the cryogenic (green) and room temperature structures (yellow). (**I, J**) Inset views of Tb-complex and Gd-complex, shown in magenta-colored sticks with metal atoms shown as grey spheres along with (2F_o_ – F_c_) map in blue, and anomalous maps in green (contoured at 7.0 sigma). PDB codes ACER3 cryo: 6YXH, ADIPOR2 cryo: 6YX9, ADIPOR2 rt: 6YXD, ADIPOR2-Tb-XO4: 6YXG, ADIPOR2-Gd-DO3: 6YXF.

**Table 1.** Crystallography data collection and refinement statistics.

| Protein | ACER3 – cryo | ADIPOR2 – cryo | ADIPOR2 – RT | ADIPOR2– Tb-XO<sub>4</sub> | ADIPOR2–Gd-DO<sub>3</sub> |
| --- | --- | --- | --- | --- | --- |
| PDB ID | 6YXH | 6YX9 | 6YXD | 6YXG | 6YXF |
| **Data Collection** |  |  |  |  |  |
| Photon energy (keV) | 12.4 | 12.4 | 12.4 | 7.5 | 7.2 |
| Beam size (μm<sup>2</sup>) | 20 x 10 | 20 x 10 | 10 x 5 | 20 x 10 | 20 x 10 |
| Synchrotron | Swiss Light Source (SLS) | Swiss Light Source (SLS) | P14 - PetraIII, Desy | Swiss Light Source (SLS) | Swiss Light Source (SLS) |
| Detector | EIGER-16M | EIGER-16M | EIGER-16M (4M ROI) | EIGER-16M | EIGER-16M |
| Photon flux (ph/s) | $6.1 \times 10^{11}$ | $6.1 \times 10^{11}$ | $6.1 \times 10^{13}$ | $6.1 \times 10^{11}$ | $6.1 \times 10^{11}$ |
| Exposure time (ms)/frame | 100 | 100 | 1.3 | 100 | 100 |
| Oscillation range / crystal | 10° | 10° | 0.6° | 10° | 10° |
| Collection strategy | 10° wedge/crystal | 10° wedge/crystal | helical line-scan on each drop; ~400 lines; 2 and 10 $\mu\text{m}$ vertical and horizontal steps respectively. | 10° wedge/crystal | 10° wedge/crystal |
| Total no. of frames | NA | NA | 382,552 | NA | NA |
| # Hits (datasets collected) | 21,250 (425) | 15,750 (315) | 28,949 | 28,500 (570) | 20,500 (410) |
| # Indexed frames† | 18,550 (371) | 5,200 (104) | 25,297 | 23,350 (467) | 17,800 (356) |
| # Datasets merged | 101 | 100 | N/A | 210 | 96 |
| # CD boluses | 9 | 1 | 5 | 16 | 7 |
| # Crystals/bolus | ~17 | ~33 | ~100 | ~27 | ~22 |
| Effective collection time† (minutes) | 8.4 (30.9) | 8.3 (26.2) | 20 (76) | 38.9 (47.5) | 29.6 (35) |
| Protein consumption ($\mu\text{L}$) | 1.6 | 0.3 | 1.5 | 4.8 | 2.1 |
| Crystal size ($\mu\text{m}^3$, approx.) | 20 x 10 x 5 | 30 x 20 x 5 | 30 x 20 x 5 | 30 x 20 x 5 | 30 x 20 x 5 |
| Space group | C222<sub>1</sub> | P2<sub>1</sub>22<sub>1</sub> | P2<sub>1</sub>22<sub>1</sub> | P2<sub>1</sub>22<sub>1</sub> | P2<sub>1</sub>22<sub>1</sub> |
| Unit-cell: a, b, c (Å) | 61.5, 69.9, 259.5 | 74.7, 100.6, 109.7 | 75.4, 100.8, 114.9 | 74.6, 100.7, 110.5 | 74.6, 100.6, 111.2 |
| **Structure refinement** |  |  |  |  |  |
| Resolution range* (Å) | 46.2 – 2.6 (2.62 – 2.6) | 22 – 2.4 (2.49 – 2.40) | 50 – 2.9 (2.98 – 2.9) | 48.0 – 3.0 (3.1 – 3.0) | 48.7 – 3.0 (3.1 – 3.0) |
| # Unique reflections* | 17,701 (1,275) | 30,556 (2,326) | 15,292 (1,971) | 32,012 (2,150) | 31,468 (2,336) |
| Multiplicity* | 36.01 (35.28) | 37.12 (33.69) | 130.57 (4.24) | 34.82 (13.14) | 14.98 (6.18) |
| CC<sub>1/2</sub>* | 0.99 (0.330) | 0.99 (0.561) | 0.98 (0.737) | 0.99 (0.3) | 0.98 (0.3) |
| I/σ(I)* | 8.4 (0.98) | 8.3 (1.02) | 8.6 (1.4) | 7.33 (1.5) | 5.06 (1.1) |
| Completeness (%) | 100 (100) | 99.4 (100) | 100 (75.7) | 99.9 (100) | 98.8 (99.7) |
| Wilson B-factor (Å<sup>2</sup>) | 65.9 | 60.7 | 42.3 | 79.1 | 78.9 |
| R-work / R-free (%) | 23.2 (32.2) / 28.0 (34.9) | 20.7 (38.0) / 24.8 (40.6) | 19.9 (23.4) / 25.3 (39.3) | 21.5 (34.6) / 25.2 (50.3) | 21.4 (30.6) / 25.9 (40.8) |
| Clashscore | 9.2 | 8.1 | 6.4 | 10.9 | 11.1 |
| RMS-bond (Å) | 0.01 | 0.005 | 0.003 | 0.01 | 0.01 |
| RMS-angle (°) | 1.06 | 0.710 | 0.659 | 1.10 | 1.15 |
| Ramachandran favoured / outlier (%) | 93.97 / 1.44 | 98.22 / 0.0 | 98.02 / 0.0 | 93.76 / 0.97 | 92.48 / 0.99 |

\* values in the parenthesis represent high-resolution shells

To demonstrate that our pipeline can also be applied to ligand discovery, we soaked ADIPOR2 crystals with a library compose of 60 small molecule fragments. We soaked each fragment into two *in meso* boluses containing ADIPOR2 microcrystals in order to ensure collection of a complete dataset. Soaked crystals were harvested, subjected to SSX diffraction data collection, data processing and ligand detection (**STAR Methods**). Structures were refined against the data using the fully automated Pipedream pipeline from Global Phasing, including structure refinement with autoBuster and automated ligand fitting with Rhofit (**Figure 3**). This led to the identification of two fragment hits, named B08 and F04 fragments (**Figure S5**) demonstrating the feasibility of this automated pipeline for ligand screening.

## Conclusions

The combination of *in meso* crystallization with CrystalDirect technology enables facile *in situ* data collection and automation of *in meso* crystal soaking, harvesting and processing producing high-resolution membrane protein structures (**Figure 4**). This approach removes critical bottlenecks in sample handling and streamlines SSX diffraction analysis of membrane proteins. In combination with CRIMS, providing web access to experimental design, analysis and crystal mounting interfaces, as well as an automated SSX data processing pipeline, we provide for the first time a fully automated, protein-to-structure pipeline for the crystallographic analysis of membrane proteins remotely operated over the web. The method is versatile, enabling structure determination both at room temperature and cryogenic conditions from a single crystallization support. All workflows described are currently available at the HTX platform at EMBL Grenoble (https://htxlab.embl.fr/) and can be reproduced at other laboratories, as the technology is readily available. The technique is compatible with standard beamline set ups, thereby eliminating the need to adapt crystal production protocols to the type of experiment or sample delivery system. This approach facilitates the systematic analysis of membrane proteins at different temperatures helping to identify conformational flexibility in membrane proteins and contributing to study receptor dynamics (Fraser et al., 2011). We have demonstrated this by capturing aspects of intramembrane enzyme:product dynamics. Finally, *in meso* ligand soaking experiments are key to support structure guided drug design against membrane proteins (Ishchenko et al., 2019; Rucktooa et al., 2018). The approach presented here uniquely combines a web-based CRIMS database and CrystalDirect platform providing a complete automated workflow for high throughput membrane protein ligand discovery.

## Acknowledgements

We want to thank Thomas Schneider and the EMBL Hamburg Team for excellent support with data collection at P14 of the PetraIII synchrotron (DESY, Hamburg, Germany), the staff at the Swiss Light Source (Paul Scherrer Institut, Villigen, Switzerland) as well as Frank Felisaz and the EMBL Grenoble Diffraction Instrumentation Team for support during the design of the plate adaptor. This work benefited from access to the HTX Lab at EMBL Grenoble. This project was supported by funding from the European CommunityH2020 Programme under the projects iNEXT (Grant No 653706) and iNEXT Discovery (Grant No 871037) as well as the Région Auvergne-Rhône-Alpes through the Booster programme. This research has received funding from the European Union’s Horizon 2020 research and innovation program under grant agreement n.° 730872, project CALIPSOplus. This project has also received funding from the European Research Council (ERC) under the European Union’s Horizon 2020 research and innovation program (grant agreement N° 647687).

## Author Contributions

RH, SB and CL produced the samples, and carried out serial data collection and structural analysis. ASH, FD and AP carried out crystallization experiments, adapted the CrystalDirect technology to LCP and contributed to serial data collection experiments. XC and JB performed molecular dynamics simulations. JAM and SG supervised the work and wrote the manuscript with RH and SB. All authors contributed to the final version of the manuscript.

## STAR Methods

### KEY RESOURCES TABLE

STAR Methods KEY RESOURCES TABLE
| REAGENT or RESOURCE | SOURCE | IDENTIFIER |
| --- | --- | --- |
| Bacterial and virus strains |  |  |
| *E. coli* cells DH5α | NEB | C2987 |
| *E. coli* cells DH10Bac | Invitrogen | 10361012 |
| Chemicals, peptides, and recombinant proteins |  |  |
| pFastBac vector | Thermofisher | 10712024 |
| pCDNA3.1 vector | ThermoFisher | V79020 |
| pMT vector | ThermoFisher | V413020 |
| Iodoacetamide | Sigma | I6125 |
| Leupetin | Euromedex | SP-04-2217 |
| Benzamidine | Sigma | B6506 |
| Phenylmethanesulfonyl fluoride (PMSF) | Sigma | P7626 |
| *n*-dodecyl-beta-D-maltopyranoside (DDM) | Anatrace | D310 |
| Lauryl maltose neopentyl glycol (MNG) | Anatrace | NG310 |
| Cholesterol hemisuccinate (CHS) | Sigma | C6512 |
| ANTI-FLAG® M2 Agarose Affinity Gel | Sigma-Aldrich | A2220 |
| FLAG peptide | Covalab | https://www.covalab.com/peptide-synthesis |
| EX-CELL420 Serum-Free Medium | Sigma-Aldrich | 14420 |
| Strep-Tactin® Superflow® high capacity resin | IBA | 2-1206 |
| Desthiobiotin | IBA | 2-1000 |
| Puromycin dihydrochloride | ThermoFisher | A1113803 |
| Transfection reagent Sf9 (Transfectin) | BioRad | 1703350 |
| Transfection reagent S2 (Effectene) | Qiagen | 301425 |
| Cadmium chloride hydrate | Sigma | 208299 |
| **Experimental models: cell lines** |  |  |
| insect cell line Sf9 | Life Technologies | 11496015 |
| insect cell line S2 | ThermoFisher | R69007 |
| **Recombinant DNA** |  |  |
| pMT-scFv-ADIPOR | Vasiliauskaitė-Brooks et al., 2017 | N/A |
| pFastBac-ADIPOR2(100-386) | Vasiliauskaitė-Brooks et al., 2017 | N/A |
| pFastBac-ACER3(1-244)-BRIL | Vasiliauskaitė-Brooks et al., 2018 | N/A |
| **Deposited data** |  |  |
| Cryogenic structure of ADIPOR2 at 2.4 Å ligand determined by SSX crystallography using CrystalDirect | This paper | PDB: 6YX9 |
| Cryogenic structure of ACER3 at 2.6 Å determined by SSX crystallography using CrystalDirect | This paper | PDB: 6YXH |
| Room temperature structure of ADIPOR2 at 2.9 Å ligand determined by SSX crystallography using CrystalDirect | This paper | PDB: 6YXD |
| Cryogenic structure of ADIPOR2 with Gd-DO3 ligand determined by SSX crystallography using CrystalDirect | This paper | PDB: 6YXF |
| Cryogenic structure of ADIPOR2 with Tb-XO4 ligand determined by SSX crystallography using CrystalDirect | This paper | PDB: 6YXG |
| **Software and algorithms** |  |  |
| CRIMS | This paper | https://htxlab.embl.fr/ |
| PyMOL | Schrödinger | https://pymol.org/2/ |
| XDS | Kabsch, 2010 | https://xds.mr.mpg.de/ |
| CrystFEL | White et al., 2012 | https://www.desy.de/~twhite/crystfel/ |
| sxdm | Basu et al., 2019 | <a href="https://github.com:shibom/sxdm.git">git@github.com:shibom/sxdm.git</a> |
| CCP4 | doi: 10.1107/S0907444910045749 | http://www.ccp4.ac.uk/ |
| COOT | Emsley and Cowtan, 2004 | https://www2.mrc-lmb.cam.ac.uk/personal/pemsley/coot/ |
| Buster | Vonrhein et al., 2006 | http://www.globalphasing.com/buster/ |
| Pipedream | Sharff et al., 2011 | https://www.globalphasing.com/buster/manual/pipedream/manual/index.html#_pipedream |
| H++ server | Gordon et al., 2005 | http://biophysics.cs.vt.edu/ |
| GROMACS 5.1 | Van Der Spoel et al., 2005 | https://www.gromacs.org |
| PLUMED 2.3 | Tribello et al., 2014 | https://www.plumed.org/ |
| PACKMOL-Memgen | Schott-Verdugo and Gohlke, 2019 | https://github.com/alanwilter/acpype/blob/master/amber19-0_linux/bin/packmol-memgen |

### RESOURCE AVAILABILITY

#### Lead contacts

Further information and requests for reagents should be directed to and will be fulfilled by the lead contacts, Sébastien Granier and José Antonio Márquez.

#### Materials availability

DNA constructs generated by the authors can be obtained upon request from the Lead Contact, but we may require a payment and/or a completed Materials Transfer Agreement if there is potential for commercial application.

#### Data and code availability

The article includes all datasets generated or analyzed during this study.

### EXPERIMENTAL MODEL AND SUBJECT DETAILS

Human ADIPOR2 and ACER3 were expressed in Sf9 cells infected with recombinant baculovirus (pFastBac, Invitrogen). Synthetic anti-ADIPOR scFv was expressed in S2 cells transfected with plasmid DNA (pMT, ThermoFisher).

### QUANTIFICATION AND STATISTICAL ANALYSIS

Quantification and statistical analyses of data are described in Method Details and Figure legends.

### METHOD DETAILS

#### scFv Anti-ADIPOR2 expression and purification

The DNA sequence of anti-ADIPOR was subcloned into a pMT expression vector (Invitrogen) containing an N-terminal BiP secretion signal and C-terminal twin-strep tag. S2 Schneider cells (ThermoFisher) expressing the scFv were induced for 10 days with cadmium chloride and supernatant containing scFv was harvested. The supernatant was concentrated and loaded onto a StrepTactin column (IBA) in scFv wash buffer (100 mM Tris pH 8, 150 mM NaCl, 1 mM EDTA), the column was washed then scFv was eluted using scFv wash buffer containing 2.5 mM desthiobiotin (IBA). The eluted scFv was concentrated and injected on a Superdex S200 size exclusion chromatography column in SEC scFv buffer (10 mM HEPES pH 7.5, 150 mM NaCl). Eluted fractions were concentrated to 15 mg/mL, snap-frozen in liquid nitrogen prior to complexation with purified receptor.

#### ADIPOR2(100-386) expression, purification and complexation with scFv

We utilized an N-terminally truncated human ADIPOR2 construct containing an N-terminal Flag-tag for affinity purification. Recombinant baculoviruses were generated using the pFastBac baculovirus system (ThermoFisher). High titer baculovirus encoding ADIPOR2 gene were used to infect *Sf9* cells at a cell density of 4 × 10^6^ cells per ml in suspension in EX-Cell 420 serum free media (Expression System). Cells were harvested by centrifugation 48 h post-infection and stored at −80 °C until purification.

Cell pellets were resuspended in lysis buffer (10 mM Tris-HCl pH 7.5, 1 mM EDTA buffer containing 2 mg·mL^-1^ iodoacetamide and protease inhibitors) without salt to lyse the cells by hypotonic lysis. Lysed cells were centrifuged (38,420 g) and the membranes were solubilized for 1 hour at 4 degrees in solubilization buffer (20 mM HEPES pH 7.5, 100mM NaCl, 1% (w/v) n-dodecyl-β-D-maltoside (DDM), 0.1% (w/v) cholesteryl hemisuccinate (CHS), 2 mg·mL^-1^ iodoacetamide and protease inhibitors). The solubilized receptor was recovered by centrifugation (38,420 g), loaded onto anti-Flag M2 column (Sigma) and washed thoroughly with DDM wash buffer (20 mM HEPES pH 7.5, 200 mM NaCl, 0.025% (w/v) DDM and 0.0001% (w/v) CHS). While on the M2 antibody resin, the detergent was exchanged to lauryl maltose neopentyl glycol (MNG, Anatrace) using MNG exchange buffer (20 mM HEPES pH 7.5, 100 mM NaCl, 0.5% (w/v) MNG). The detergent exchange was performed by washing the column with a series of seven buffers (3 CV each) made up of the following ratios (%/%) DDM wash buffer and MNG exchange buffer: 100:0, 50:50, 25:75, 10:90, 5:95, 1:99, 0:100. The column was then washed with MNG wash buffer (20 mM HEPES pH 7.5, 100 mM NaCl, 0.02% (w/v) MNG) and the bound receptor was eluted in the same buffer supplemented with 0.2 mg·mL^-1^ Flag peptide (Covalab).

Eluted ADIPOR2 was incubated with purified antiADIPOR-scFv at a molar ratio of 2:1, ADIPOR2:scFv on ice for 30 minutes to form a complex, then loaded onto a StrepTactin column (IBA). The complex was washed with MNG wash buffer and the bound complex was eluted in the same buffer supplemented with 2.5 mM desthiobiotin. Eluted complex was concentrated to 500 μL and injected on a Superdex S200 size exclusion chromatography column in SEC MNG buffer (20 mM HEPES pH 7.5, 100 mM NaCl, 0.002% (w/v) MNG). Purified complex was concentrated to 15 mg/mL and snap-frozen in liquid nitrogen prior to crystallization.

#### ACER3(1-244) expression and purification

We utilized a C-terminally truncated human ACER3 construct containing an N-terminal Flag-tag for affinity purification. Recombinant baculoviruses were generated using the pFastBac baculovirus system (ThermoFisher). High titer baculovirus encoding ACER3 gene were used to infect *Sf9* cells at a cell density of 4 × 10^6^ cells per ml in suspension in EX-Cell 420 serum free media (Expression System). Cells were harvested by centrifugation 48 h post-infection and stored at −80 °C until purification.

Cell pellets were resuspended in lysis buffer (10 mM Tris-HCl pH 7.5, 1 mM EDTA buffer containing 2 mg·mL^-1^ iodoacetamide and protease inhibitors) without salt to lyse the cells by hypotonic lysis. Lysed cells were centrifuged (38,420 g) and the membranes were solubilized for 1 hour at 4 degrees in solubilization buffer (20 mM HEPES pH 7.5, 100mM NaCl, 1% (w/v) n-dodecyl-β-D-maltoside (DDM), 0.1% (w/v) cholesteryl hemisuccinate (CHS), 2 mg·mL^-1^ iodoacetamide and protease inhibitors). The solubilized enzyme was recovered by centrifugation (38,420 g), loaded onto anti-Flag M2 column (Sigma) and washed thoroughly with DDM wash buffer (20 mM HEPES pH 7.5, 200 mM NaCl, 0.025% (w/v) DDM and 0.0001% (w/v) CHS). While on the M2 antibody resin, the detergent was exchanged to lauryl-maltose-neopentyl-glycol (MNG, Anatrace) using MNG exchange buffer (20 mM HEPES pH 7.5, 100 mM NaCl, 0.5% (w/v) MNG). The detergent exchange was performed by washing the column with a series of seven buffers (3 CV each) made up of the following ratios (%/%) DDM wash buffer and MNG exchange buffer: 100:0, 50:50, 25:75, 10:90, 5:95, 1:99, 0:100. The column was then washed with MNG wash buffer (20 mM HEPES pH 7.5, 100 mM NaCl, 0.02% (w/v) MNG) and the bound enzyme was eluted in the same buffer supplemented with 0.2 mg·mL^-1^ Flag peptide (Covalab). Eluted enzyme was concentrated to 500 μL and injected on a Superdex S200 size exclusion chromatography column in SEC MNG buffer (20 mM HEPES pH 7.5, 100 mM NaCl, 0.002% (w/v) MNG). Purified ACER3 was concentrated to 18 mg/mL and snap-frozen in liquid nitrogen prior to crystallization.

#### LCP crystallization in CrystalDirect plates

Protein constructs, expression and purification were as previously described for ADIPOR2 (Vasiliauskaité-Brooks et al., 2017) and ACER3(Vasiliauskaité-Brooks et al., 2018). Snap-frozen protein aliquots were sent on dry ice to the high-throughput crystallization lab (HTX Lab) at EMBL Grenoble for crystallization set-up (Dimasi et al., 2007). Sample mesophases of ADIPOR2 and ACER3 were generated using the LCP method (Caffrey, 2015; Landau and Rosenbusch, 1996; Liu and Cherezov, 2011). Concentrated protein solutions (ADIPOR2: 15 mg/mL, ACER3: 18 mg/mL) were thawed and reconstituted with a 10:1 monoolein:cholesterol (Sigma) mixture in a ratio of 2:3 protein:lipid (v/v). The mesophase was formed by using the coupled two-syringe method (Caffrey, 2015; Caffrey and Cherezov, 2009; Liu and Cherezov, 2011). The mesophase was dispensed onto the crystallization film of CrystalDirect plates (MiTeGen, ref. M-XDIR-96-2) in 30 or 50 nl bolus and overlaid with 600 nl precipitant solution using either a Mosquito LCP robot (SPT Labtech) equipped with humidifier or a Gryphon LCP robot (Art Robbins Instruments). Reservoir wells next to each mesophase bolus contained 45 μL of precipitant solution to maintain drop humidity during crystallization (**Figure 3**). Precipitant solutions contained 42.3% PEG 400, 110 mM potassium citrate, 100 mM HEPES pH 7.0 for ADIPOR2 and 39.5–41% PEG 400, 100□mM HEPES pH 7.5, 75 mM magnesium sulphate and 5% DMSO for ACER3. Under these conditions ADIPOR2 crystals were stable for several weeks.

The plates were incubated at 20°C and monitored with a Rock Imager-1000 robot (Formulatrix). In our experience the surface of the bolus in contact with the precipitant solution presents a rough structure, which interferes with crystal imaging. To overcome this, and facilitate crystal detection, a plate adaptor was developed to invert the plate inside the imager (**Figure S6)**. This adaptor consists of a plastic frame compatible with the SBS-plate format into which the crystallization plate is inserted and turned upside-down (crystallization film facing up) whilst guaranteeing safe handling by the imaging robot. Thanks to this adaptor; crystallization experiments are imaged through the film in which the LCP bolus is resting, therefore presenting a smooth and flattened surface to the camera, notably improving crystal imaging.

Design and follow-up of crystallization experiments was carried out remotely using CRIMS. Crystals were observed within one day and reached full size within five days: ~30□×□20□x □5□ μm^3^ for ADIPOR2 in cubic phase and ~20□x□10□x□5□μm^3^ for ACER3 in sponge phase (**Figure S4**).

CrystalDirect plates are compatible with standard crystallization robotics, which makes setting LCP crystallization experiments very straightforward. They can be used both for crystallization screening and optimization. In our experience it was straightforward to transfer LCP crystallization from sandwich to CrystalDirect plates and vice versa by minor adjustment of the precipitant concentration.

#### Room temperature serial crystallography data collection

*In situ* diffraction experiments were carried out at the P14 beam line operated by EMBL Hamburg at the PETRA III storage ring (Gati et al., 2014) (DESY, Hamburg, Germany). While the crystallization film used in CrystalDirect plates is extremely thin (20 μm), the plate is typically sealed on the top side with a standard film (c.a. 100 μm thick). In order to reduce the scattering background, the top sealing film was removed from the plates and replaced with a CrystalDirect (Zander et al., 2016) *in situ* plate seal (Mitegen, ML-CDSF1-10, with the same thickness as the crystallization film) immediately before data collection. The CrystalDirect plates were mounted on a SBS-plate goniometer. Data collection was carried out by rastering each LCP bolus applying multiple vertical “helical-lines” separated by 10 μm in the horizontal direction and applying a continuous oscillation with the plate goniometer from the bottom to the top of the line (**Supplementary Video 2**). A 10 x 5 μm^2^ beam operated at 12.4 KeV and 100% transmission at a photon flux of 1.17 x 10^13^ was used, which delivered roughly 100 kGy per frame (Gati et al., 2014). The length and number of lines as well as the oscillation were adjusted to match the size of each bolus. Typically, between 300 and 400 helical lines about 1.8 mm in length were necessary to sample a full LCP bolus with an oscillation of between 0.3 and 0.4 degrees per frame. At the P14 beam line optimal data collection patterns can be easily selected by simply entering the number of lines, separation and oscillation range per line in a dedicated user interface. Images were collected using an Eiger X 16M operated in 4M mode compatible with a fast exposure time of 1.3 msec. A typical data collection session took four minutes per bolus (including focusing and centering the bolus into the beam). For ADIPOR2 five LCP boluses and about 20 minutes of data collection were sufficient to obtain a full dataset. This consisted of 382,552 diffraction images, out of which 28,949 were crystal hits and 25,297 diffraction images were indexable (**Table 1**).

#### Automated, laser-based LCP-crystal harvesting and cryocooling

Automated harvesting of crystals grown in mesophase media was carried out with the help of the CRIMS software and the CrystalDirect harvester robot (Cipriani et al., 2012; Márquez and Cipriani, 2014; Zander et al., 2016) (**Supplemental Video 1**). Briefly, crystallization images taken by the crystal imaging farm are presented through a CRIMS web interface (**Figure S2**). This interface enables scientists to select from a collection of different shapes that can then be placed onto the crystallization image and rotated for optimal crystal recovery. The scientist has a number of choices concerning sample mounting. For example, it is possible to mount multiple crystals in a single pin or use the laser to isolate single crystals (**Figure S1**). It is also possible to perform multiple crystal harvests from a single drop. All harvesting operations are recorded by the CRIMS system.

To harvest the samples, the crystallization plate is transferred to the CrystalDirect harvester from which point all harvesting operations are automatically performed (**Supplemental Video 1**). The harvester uses a femto-second laser beam operating in the photoablation regime enabling precise excision of the crystallization film while avoiding heat transfer or mechanical damage to the sample (Aragón et al., 2019; Bezerra et al., 2017; Martin-Malpartida et al., 2017). The harvester robot uses the plate barcode to recover all harvesting operations stored by CRIMS. A camera system along with automated image recognition algorithms allows the harvester to automatically locate crystallization experiments and precisely position the harvesting shapes according to the design specified through the CRIMS harvesting interfaces. After laser excision, the film piece containing the crystals is automatically attached to a SPINE standard X-ray data collection pin and transferred to a nitrogen gas stream at 100 K. Cryo-cooled samples are then recovered by a robotic arm that transfers them to a cryogenic sample storage dewar. The system can work both with SPINE and UNIPUCK puck formats that are compatible with most synchrotron beam lines. Multiple harvests can be performed consecutively in batch-mode. The sample storage dewar has capacity for 312 pins. All data, including sample, pin and puck locations for each sample along with harvesting images automatically generated during the harvesting processes is transferred back and stored in CRIMS.

#### Soaking of ADIPOR2 crystals

The CrystalDirect LCP crystallization method facilitates post-crystallization treatments, including soaking, by enabling the removal of the top sealing film without disturbing the bolus. Two commercially available caged-metal complexes (Engilberge et al., 2017; Girard et al., 2003) were used for soaking experiments, lanthanide phasing compound Gd-HPDO3A (Jena Bioscience PK-CSM002-0002A) at a concentration of 500 mM and polyvalan Crystallophore kit Tb-XO4 (Molecular Dimensions ref: MD2-80) at 100 mM. For soaking experiments, the top sealing film of the CrystalDirect plate was removed and 30–150 nl of Tb-XO4 or 60–120 nl of Gd-DO3 ligand solutions were delivered directly to the LCP crystallization drops with a SPT-Labtech crystallization robot, after which the plate was resealed. This process takes a few seconds for a drop and just a couple of minutes for the whole plate, avoiding dehydration of the LCP drops. Plates were resealed and crystals were soaked for 90 minutes at 20°C before crystal harvesting. The final concentrations of ligands were approximately 5–25 mM Tb-XO4 and 10–25 mM Gd-DO3.

#### Cryogenic serial crystallography data collection

Diffraction data from samples vitrified using the CrystalDirect technology were collected remotely at the X06SA beamline of the Swiss Light Source (SLS), Villigen, Switzerland. After sample transfer to the goniometer, loops containing multiple crystals were rapidly raster-scanned (**Figure S3**) with the X-ray beam to identify positions corresponding to diffracting crystals(Basu et al., 2019; Wojdyla et al., 2016). In a second step 10° wedges were collected with 0.1 sec exposure time from each of the identified positions at a speed of 2°/sec. If dose permitted, 2-3 sweeps of 10° wedges were collected from each of those identified crystals. Each 10° wedge takes typically 5 sec. Effective data collection time required to obtain a complete dataset from each target is summarized in **Table 1**. Diffraction images were collected using an EIGER X 16M detector (Dectris) with a sample-to-detector distance of 30 cm, beam energy was 12.4 KeV for apo crystals, and 7.2 KeV for soaked crystals in order to exploit the anomalous signals from the heavy elements. The beam size was 10×20 μm^2^ and 100% transmission.

#### Data processing and structure refinement

Initial diffraction patterns, collected under cryogenic condition at the SLS, were processed using XDS (Kabsch, 2010), followed by real-time automated selection and merging of statistically equivalent datasets with XSCALE using the *sxdm* tool (Basu et al., 2019) as deployed at the beamline. Datasets were selected using a criterion or a combination of multiple criteria, including ISa cut-off of 3.0, unit-cell, pair-CC based hierarchical clustering as well as Δ-CC_1/2_ iteratively (Assmann et al., 2020). For *in situ* datasets collected at PetraIII P14 beamline, we used an in-house script to run CrystFEL (White et al., 2012) version 0.8.0 automatically as the data had arrived. This allowed us to quickly identify hits and reduce data volume and produce a mtz file ready for structural analysis. All the structures were solved by molecular replacement using PHASER and previously determined structures of ACER3 and ADIPOR2 as search models (PDB IDs 6G7O, 5LWY respectively). The resolution cut-offs for all structures were determined by a combined metric of CC_1/2_ of 0.30 and I/σ(I) of 1.0 (Karplus and Diederichs, 2012). Structure refinement was carried out iteratively with BUSTER (Vonrhein et al., 2006) version 2.10.3 and Coot(Emsley and Cowtan, 2004). At late stages of refinement, translation liberation screw-motion parameters generated within BUSTER (Vonrhein et al., 2006) were used. MolProbity (Williams et al., 2018) was used to assess the quality of the structure and the data collection and refinement statistics are summarized in **Table 1**. The data refinement and analysis software were compiled and supported by the SBGrid consortium (https://sbgrid.org/).

##### ADIPOR2-RT

About 3.8 million diffraction patterns were collected from 19 drops in 1 hour and 16 minutes approximately from 19 LCP boluses in one 96-well plate, out of which 170,944 images correspond to crystal hits. However, a complete room-temperature dataset was obtained from only 5 drops in a 96 well-plate, which yielded 25,297 indexed diffraction patterns. The room-temperature structure was refined at 2.9-resolution using AUTOBUSTER with Rwork of 19.9 % and Rfree of 25.3 %. The relative lower resolution for ADIPOR2-RT over ADIPOR2-Cryo is owing to the 10-fold less deposited dose at room-temperature. The structure was determined from images obtained from 5 drops, requiring 20 min of collection time.

##### ADIPOR2-Cryo

315 sweeps of 10° wedges were collected from only 1 loop, containing about 33 microcrystals. 104 datasets were successfully indexed and processed using XDS, out of which 100 datasets were selected based on ISa value cutoff of 3.0, followed by merging with XSCALE. The final mtz file, obtained from automated pipeline, was used for structure determination and refined at 2.4 Å resolution using AUTOBUSTER with Rwork of 20.7 % and Rfree of 24.8 %. The structure was determined from 100 datasets, obtained from 8.3 min of collection time.

##### ACER3-Cryo

425 sweeps of 10° wedges were collected from 9 CrystalDirect loops, out of which 375 datasets were successfully processed. 101 datasets were selected based on Δ-CC_1/2_(Assmann et al., 2020) and ISa values as implemented in *sxdm* tool (Basu et al., 2019), followed by merging with XSCALE. The final mtz file was used for structure determination in Molecular replacement method. ACER3 structure was refined at 2.6 Å resolution using AUTOBUSTER with Rwork of 23.5 % and Rfree of 28.9 %. The structure was determined from 101 datasets, obtained from 8.4 min of collection time.

##### ADIPOR2-TbXO_4_

570 sweeps of 10° wedges were collected from 16 loops, out of which 467 datasets were successfully processed in XDS, followed by 210 datasets being selected and merged with and *sxdm,* at X06SA beamline of the SLS. The Tb atoms from the soaked cluster produced strong anomalous signals. The minimum datasets required to extract sufficient anomalous signals to validate the presence of soaked ligand, were obtained in 17.5 min of collection time. The structure was refined at 3.0-resolution using AUTOBUSTER with R-work of 21.5% and Rfree of 25.2%.

##### ADIPOR2-GdDO_3_

410 x 10° wedges were collected from 7 loops, out of which 356 datasets were processed in XDS, followed by 96 datasets, being selected and merged with XSCALE automatically by *sxdm* tool, as installed at X06SA beamline of the SLS. The soaked heavy metal Gd-cluster produced a strong anomalous signal. The minimum datasets required to obtain sufficient anomalous signals to validate the presence of soaked ligand, were collected in 8 min. The structure was refined at 3.0 Å resolution using AUTOBUSTER with R-work of 21.4% and Rfree of 25.9%.

#### Molecular dynamics simulations

The initial receptor models were based on the X-ray crystal structures of ADIPOR2 in complex with an oleic acid. The protonation states of titratable residues were predicted at pH 6.5 using the H++ server (Gordon et al., 2005). The receptor systems were embedded in explicit POPC membrane and aqueous solvent neutralized with Cl^-^ ions. Effective point charges of the ligands were obtained by RESP fitting (Wang et al., 2000) of the electrostatic potentials calculated with the HF/6-31G* basis set using Gaussian 09 (Frisch et al., 2016). The Amber 99SB-ildn (Wang et al., 2004) and GAFF (Lindorff-Larsen et al., 2010) force fields were used for the protein and the ligands, respectively. Parameters for the active site were adopted from the Zinc AMBER Force Field (Peters et al., 2010). The TIP3P (Jorgensen et al., 1983) and the Joung-Cheatham models (Joung and Cheatham, 2008) were used for the water and the ions, respectively. After energy minimization, all-atom MD simulations were carried out using Gromacs 2018 (Van Der Spoel et al., 2005) patched with the PLUMED 2.3 plugin(Tribello et al., 2014). Each system was gradually heated to 298 K and pre-equilibrated during 10 ns of brute-force MD in the *NRT*-ensemble. The replica exchange with solute scaling (REST2) (Wang et al., 2011) technique was then employed to enhance the sampling with 48 replicas in the *NVT* ensemble. The protein and the ligands were considered as “solute” in the REST2 scheme-force constants of their van der Waals, electrostatic and dihedral terms were subject to scaling. The effective temperatures used for generating the REST2 scaling factors ranged from 298 K to 700 K, following a distribution calculated with the Patriksson-van der Spoel approach (Patriksson and van der Spoel, 2008). Exchange between replicas was attempted every 1000 simulation steps. This setup resulted in an average exchange probability of ~40% during 40 ns (× 48 replicas) of simulations. The first 10 ns were discarded for equilibration. Only the original unscaled replica (at 298 K effective temperature) was collected and analyzed.

