## Supplementary material for "An automated platform for structural analysis of membrane proteins through serial crystallography": Supplemmental material

### Supplemental Information

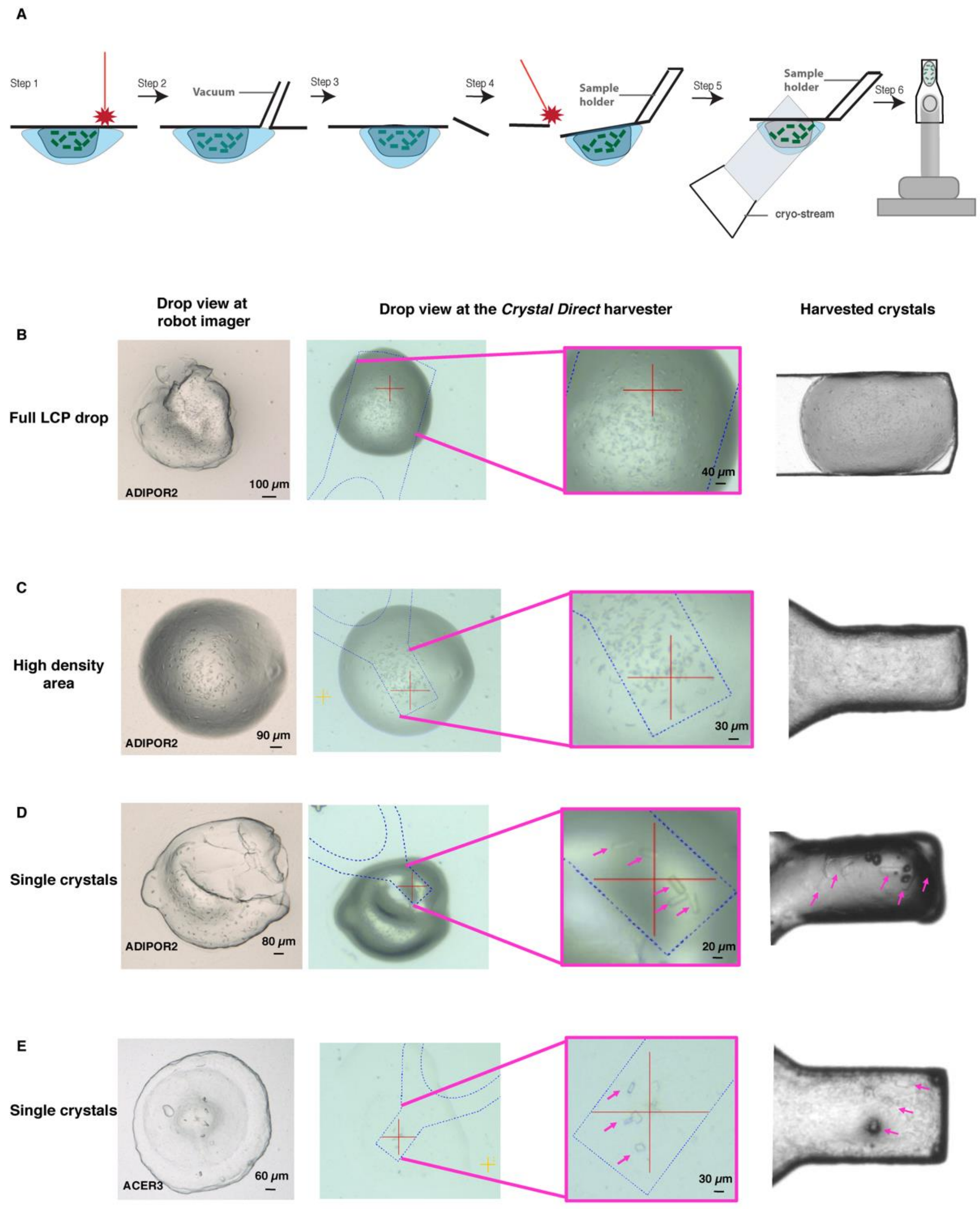


**Figure S2.** (**A**) Schematic representation of the CrystalDirect harvesting process: the laser cuts an aperture in the drop and the robot gently removes the excess of precipitant solution by aspiration, then the laser-based film excision is applied and the film piece containing the sample is automatically glued to a data collection pin and cryo-cooled. Samples are then transferred to a high-capacity cryogenic storage system (not shown). (**B–D**) Examples of ADIPOR2 and ACER3 micro-crystals grown and harvested from CrystalDirect plates. Different scenarios involving harvesting of full boluses or sections of them are presented.

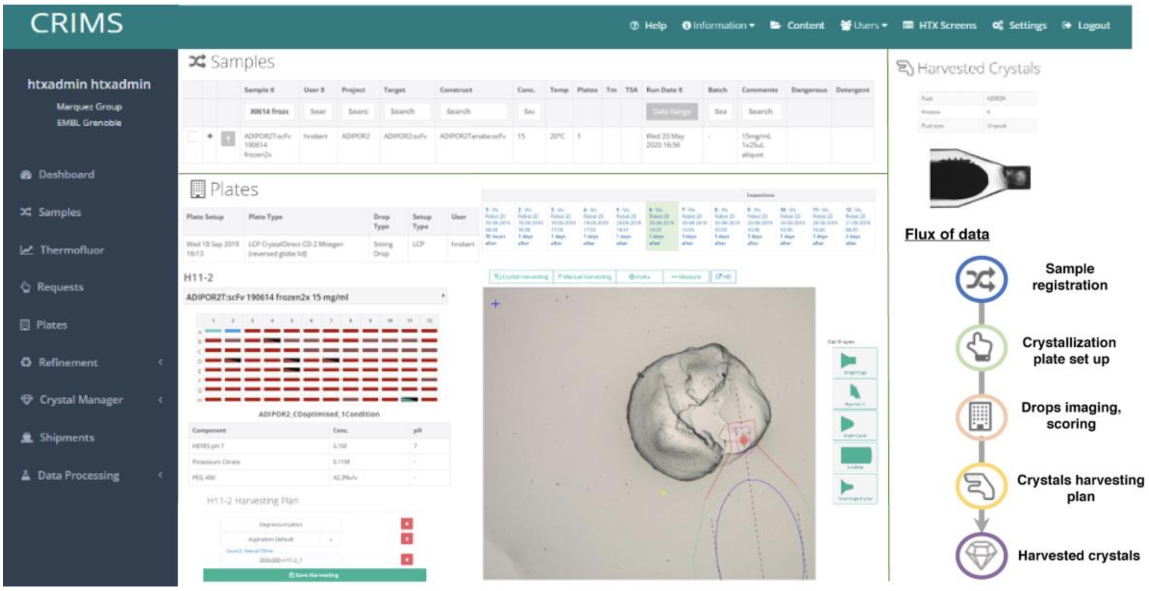


**Figure S2.** The Crystallographic Infomatnion Management System (CRIMS) provides automated sample tracking and data management over the whole experimental workflow as illustrated in the right panel. The main panel shows the crystal harvesting interface with a harvesting plan for well H11 of a CrystalDirect plate containing ADIPOR2 crystals.

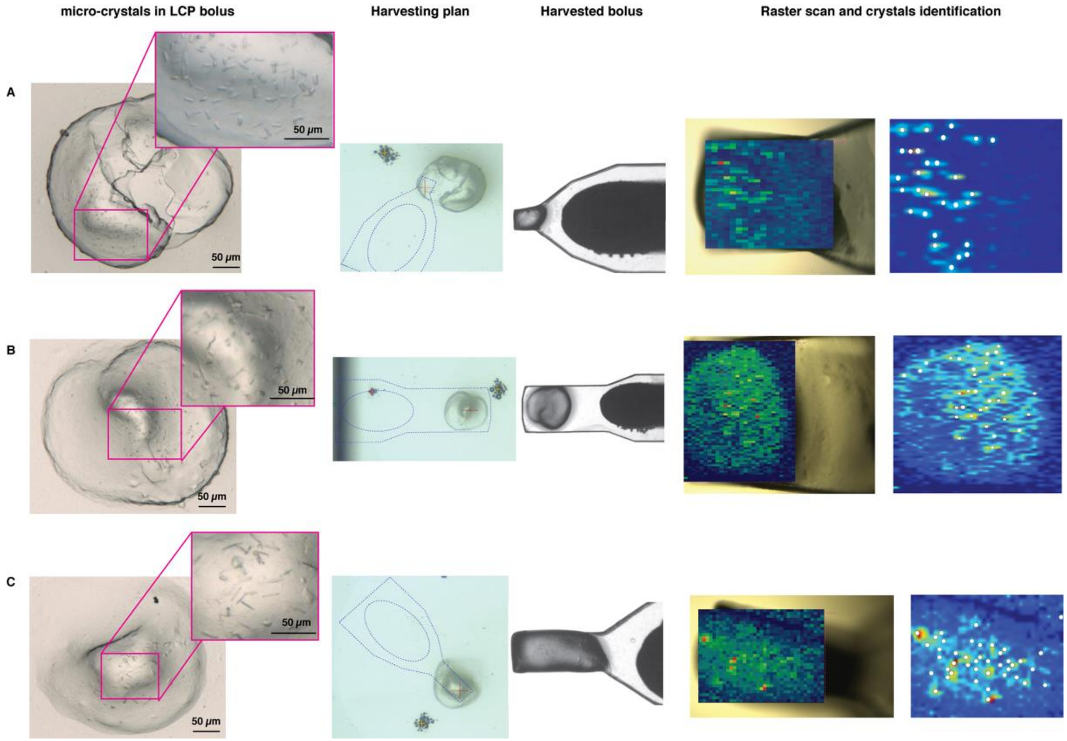


**Figure S3:** Automated harvesting and rastering of LCP crystals. Crystals grown in mesophase using a CrystalDirect plate. The panels (from left to right) illustrate different steps of the process: First column shows the microcrystals of different membrane proteins in LCP bolus; second column shows automated harvesting plan designed in CRIMS; third column shows samples harvested and cryo-cooled with the CrystalDirect robot; and final column shows the samples at the beam line with the result of the X-ray raster scan superposed at cryogenic temperature. Three representative samples – ADIPOR2, ACER3, and a GPCR target for screening processes involving harvesting of the full bolus or parts of it are shown in **A–C**, respectively.

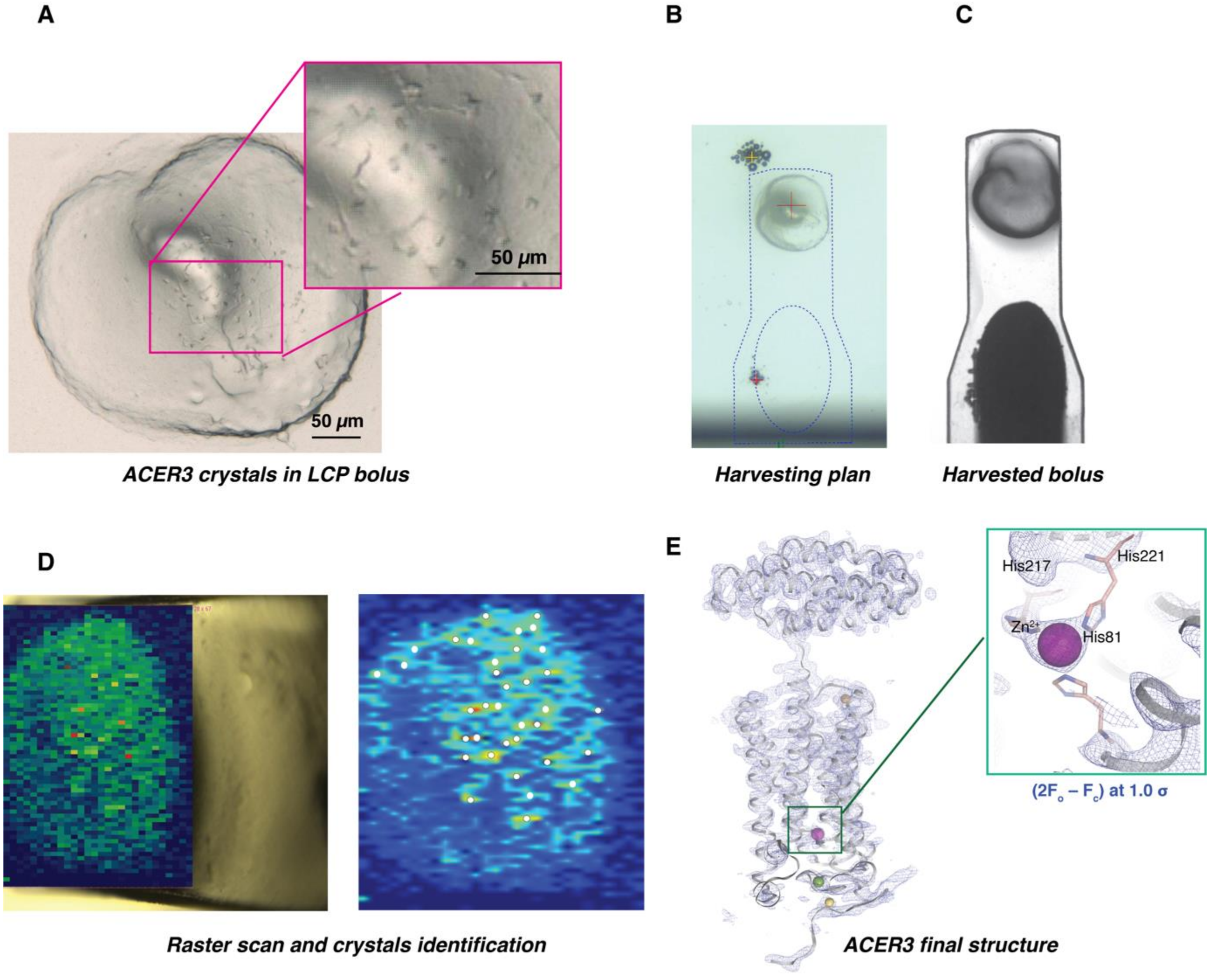


**Figure S4.** ACER3 structure determination on CrystalDirect harvested micro-crystals. (**A**) ACER3 micro-crystals. (**B**) Image from CRIMS of crystal harvesting plan and (**C**) image from CrystalDirect harvester of harvested crystals in the cryo-steam. (**D**) X-ray diffraction heat map and crystal picking for miniset collection at beamline X06SA (PXI, Swiss Light Source). (**E**) Refined structure and electron density map of ACER3 in this work (PDB: 6YXH).

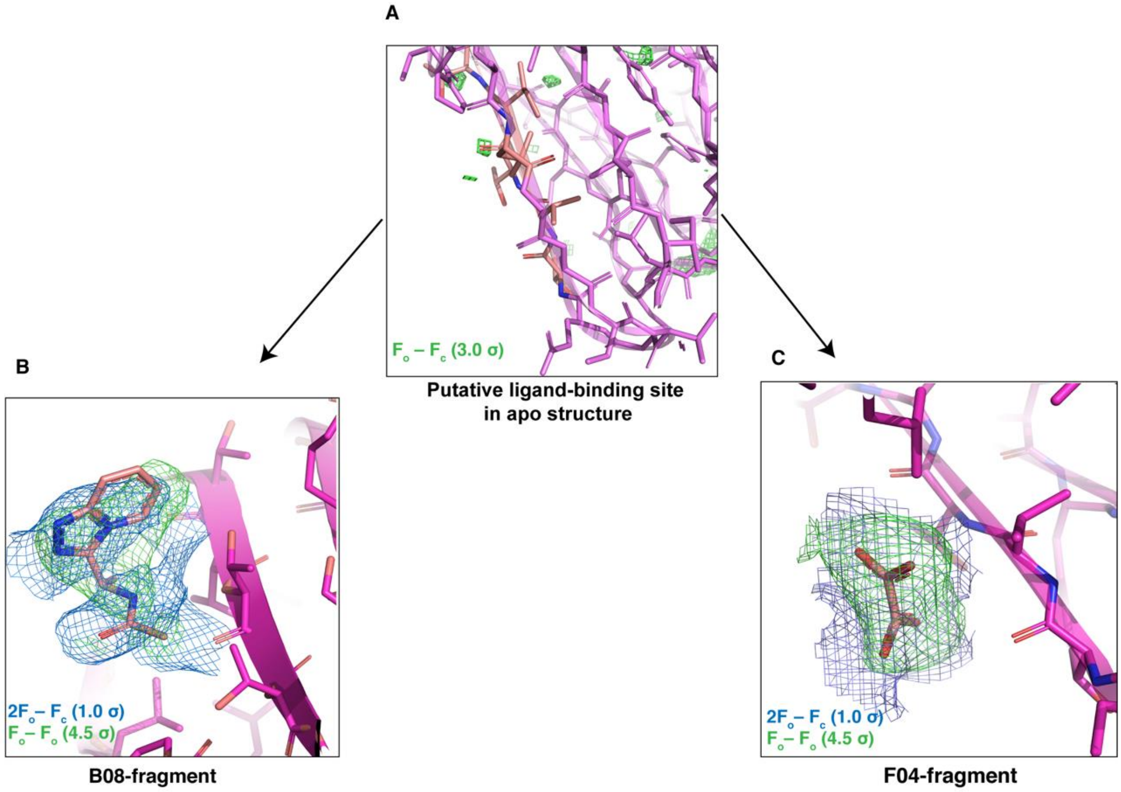


**Figure S5.** Electron density data for fragment-soaked ADIPOR2 crystals. Fo–Fc difference map (green) contoured at 3 sigma (**A**) unsoaked ADIPOR2 crystals. 2Fo–Fc in blue at 1.0 sigma and Fo–Fo in green at 4.5 sigma for soaked fragments B08 and F04 respectively, depicted as stick models (**B**, **C**).

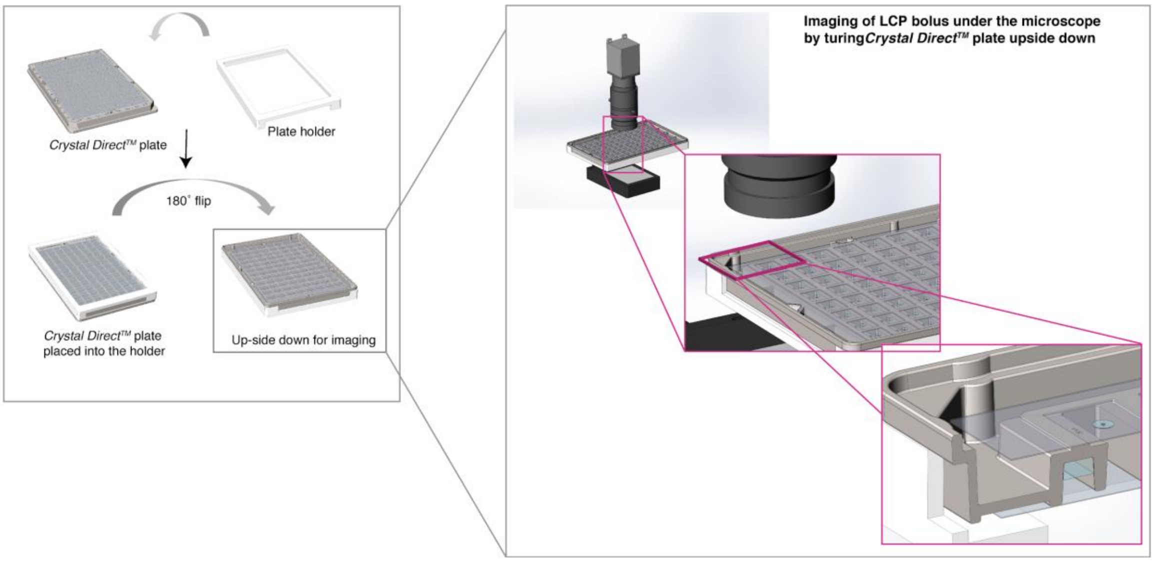


**Figure S6.** 96-well plate adaptor enabling optimal imaging of LCP experiments with commercial imaging robots. Most commercial imaging systems image crystallization plates from the top. However, the top side of the LCP bolus in CrystalDirect plates tends to present a rough structure, which may interfere with crystal imaging. To overcome this and facilitate crystal detection a plate adaptor (plate holder) was developed enabling an inverted configuration during imaging. This adaptor consists of a plastic frame compatible with the SBS-plate format into which the crystallization plate can be inserted (**left panel**) and turned upside-down (crystallization film facing up and towards the camera) whilst guaranteeing safe handling by the imaging robot (**right panel**). Thanks to this adaptor, crystallization experiments are imaged through the film in which the LCP bolus is resting, therefore always presenting a smooth and flat surface to the camera, which notably improves crystal imaging.
